# INTERACTION OF THE PERIPLASMIC CHAPERONE SURA WITH THE INNER MEMBRANE PROTEIN SECRETION (SEC) MACHINERY

**DOI:** 10.1101/2022.09.14.507990

**Authors:** Lucy Troman, Sara Alvira, Bertram Daum, Vicki A. M. Gold, Ian Collinson

## Abstract

Gram-negative bacteria are surrounded by two protein-rich membranes with a peptidoglycan layer sandwiched between them. Together they form the envelope (or cell wall), crucial for energy production, lipid biosynthesis, structural integrity, and for protection against the physical and chemical environmental challenges. To achieve envelope biogenesis, periplasmic and outer-membrane proteins (OMPs) must be transported from the cytosol and through the inner-membrane, *via* the ubiquitous SecYEG protein-channel. Emergent proteins either fold in the periplasm or cross the peptidoglycan (PG) layer towards the outer-membrane for insertion through the β-barrel assembly machinery (BAM). Trafficking of hydrophobic proteins through the periplasm is particularly treacherous given the high protein density and the absence of energy (ATP or chemiosmotic potential). Numerous molecular chaperones assist in the prevention and recovery from aggregation, and of these SurA is known to interact with BAM, facilitating delivery to the outer-membrane. However, it is unclear how proteins emerging from the Sec-machinery are received and protected from aggregation and proteolysis prior to an interaction with SurA. Through biochemical analysis and electron microscopy we demonstrate the binding capabilities of the unoccupied and substrate-engaged SurA to the inner-membrane translocation machinery complex of SecYEG-SecDF-YidC – *aka* the holo-translocon (HTL). Supported by AlphaFold predictions, we suggest a role for periplasmic domains of SecDF in chaperone recruitment to the protein translocation exit site in SecYEG. We propose that this immediate interaction with a recruited chaperone helps to prevent aggregation and degradation of nascent envelope proteins, facilitating their safe passage to the periplasm and outer-membrane.

## INTRODUCTION

Outer membrane biogenesis in Gram-negative bacteria requires targeting of the outer membrane proteins (OMPs) to the ubiquitous Sec machinery of the inner-membrane, guided by a cleavable N-terminal signal-sequence [1]. The trans-membrane proton-motive force (PMF) and ATP hydrolysis by SecA subsequently drive translocation through the hourglass-shaped SecYEG channel into the periplasm [2–4], reviewed by [5,6]. OMPs must then traverse the periplasm and peptidoglycan layer toward the β-barrel assembly machinery (BAM) for folding and insertion into the outer membrane [7,8], reviewed by [9].

Similar to the trafficking process to and across the inner-membrane, OMPs must remain unfolded during subsequent passage through the periplasm until they reach the outer-membrane. During their journey through this crowded environment [10,11], periplasmic chaperones such as SurA, Skp, PpiD and the protease DegP are recruited to help prevent and resolve aggregation [12]. Unlike cytosolic quality control factors, they must somehow operate in the absence of ATP, perhaps by virtue of their structural plasticity [13–16]. SurA is thought to be the dominant chaperone for outer-membrane delivery due to its known affinity for both the hydrophobic motifs characteristic of OMPs, and the BAM complex in the outer membrane [12,17]. Critically, the activity of SurA in inter-membrane transport depends on an interaction prior to aggregation, as unlike the Skp, it lacks the ability to recover aggregated substrates [18]. Thus, it is almost self-evident that SurA needs to interact with OMPs, prior to their release from the Sec machinery, to minimise the potential for aggregation in the periplasm. A kinetic analysis of the maturation and folding of the OMP LamB showed that *surA* mutants delay signal sequence cleavage; consistent with an interaction during, or very shortly after, translocation through the inner membrane [19].

Here, we investigate whether or not SurA makes any interaction with the inner-membrane translocation machinery, potentially for association of nascent OMPs. The core-translocon SecYEG does not have periplasmic domains large enough to accommodate SurA; however, the ancillary factors, SecDF and YidC, which associate to form the holo-translocon (HTL) do [20,21][9]. The work described here implicates these periplasmic domains in the recruitment of SurA to the translocation exit channel to help streamline the onward passage of proteins through the envelope.

## RESULTS

### THE SURA CHAPERONE INTERACTS WITH THE BACTERIAL TRANSLOCON

For initial investigation into any potential interaction between the holo-translocon (HTL) and the periplasmic chaperone SurA we used native PAGE. Purified HTL was pre-incubated for 30 minutes at 30°C with either unoccupied SurA or with the chaperone-substrate complex (SurA-OmpA) to allow for any complex association. The result visualised a faint high molecular weight band (higher than SurA and HTL alone) only in the presence of both HTL and SurA (vacant or OmpA bound; Figure 1a, asterisk). This band, and the corresponding molecular weight region from each sample (Figure 1a, black rectangle samples 1 - 5) were excised and applied to a second dimensional analysis by denaturing PAGE (Figure 1b). The analysis showed that the higher molecular weight band (Figure 1a, samples 4 and 5) contained both HTL, SurA and, when included in association with SurA, OmpA also. The identification of a distinct band in these samples, enriched in chaperone, suggests an association of the translocon irrespective of an associated substrate. However, additional lower molecular weight bands in the native gel, corresponding to HTL alone and smaller complexes (Figure 1a), suggest the HTL-SurA association is weak and the HTL is prone to dissociation. This is unsurprising, and consistent with the transient and dynamic nature of this interaction.

**Figure 1:**
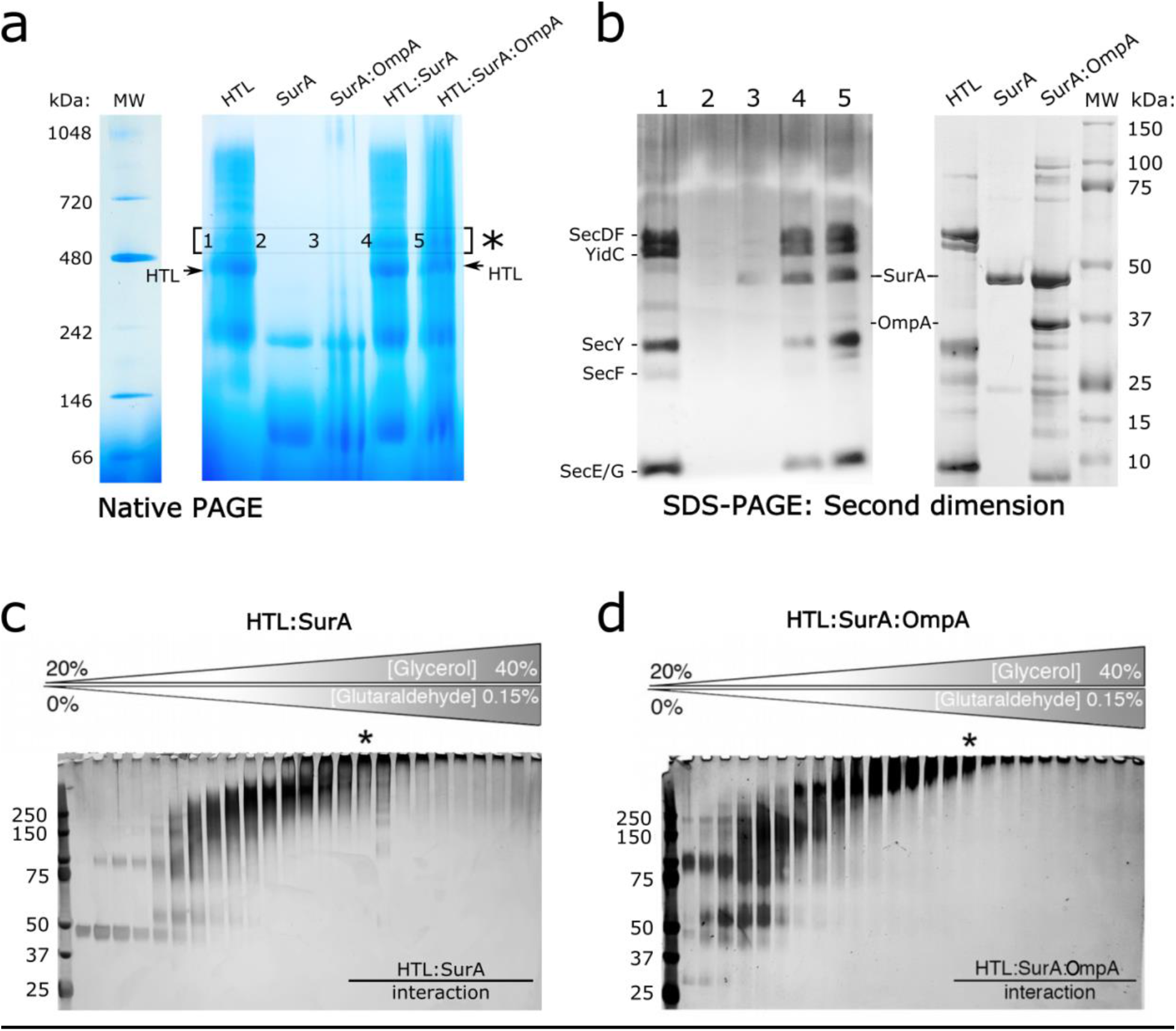
Complex formation between HTL and SurA or SurA:OmpA through NativePAGE and density gradient centrifugation: a) NativePAGE analysis of HTL, SurA, SurA:OmpA alone and mixtures of HTL:SurA and HTL:SurA:OmpA. b) Bands 1-5 from (a) denoted with an asterisk were excised at the same height and analysed by SDS-PAGE revealing the presence of protein complexes. 2 ug of HTL, SurA and SurA:OmpA were run in parallel and silver stained as markers. c,d) Silver stained SDS-PAGE analysis of glycerol fractions following crosslinking density gradient centrifugation of the translocon with SurA chaperone alone (c) or engaged with OmpA substrate (d). The asterisks in (c) and (d) denote fractions taken for further analysis via negative stain EM.

Density gradient centrifugation enables the isolation of complexes of lower-affinity interacting proteins. Additional gradient fixation, or GraFix, prevents any complex dissociation during the centrifugation through a gradient of increasing, but low, concentrations of cross-linkers [22]. We successfully applied this technique for the isolation of HTL bound to SurA and to SurA-OmpA (Figure 1c, d). The fractions were analysed by SDS-PAGE; as expected, the larger molecular weight fractions, at higher glycerol concentrations, were fully cross-linked with all the protein constituents migrating as a single band at the top of the gel. The cross-linked sample corresponding to HTL bound to SurA (Figure 1c, asterisk) was subject to mass spectroscopy, to duly confirm the presence of SurA and translocon constituents (supplementary Figure S1), followed by electron microscopy (EM; Figure 2a; supplementary Figure S2).

**Figure 2:**
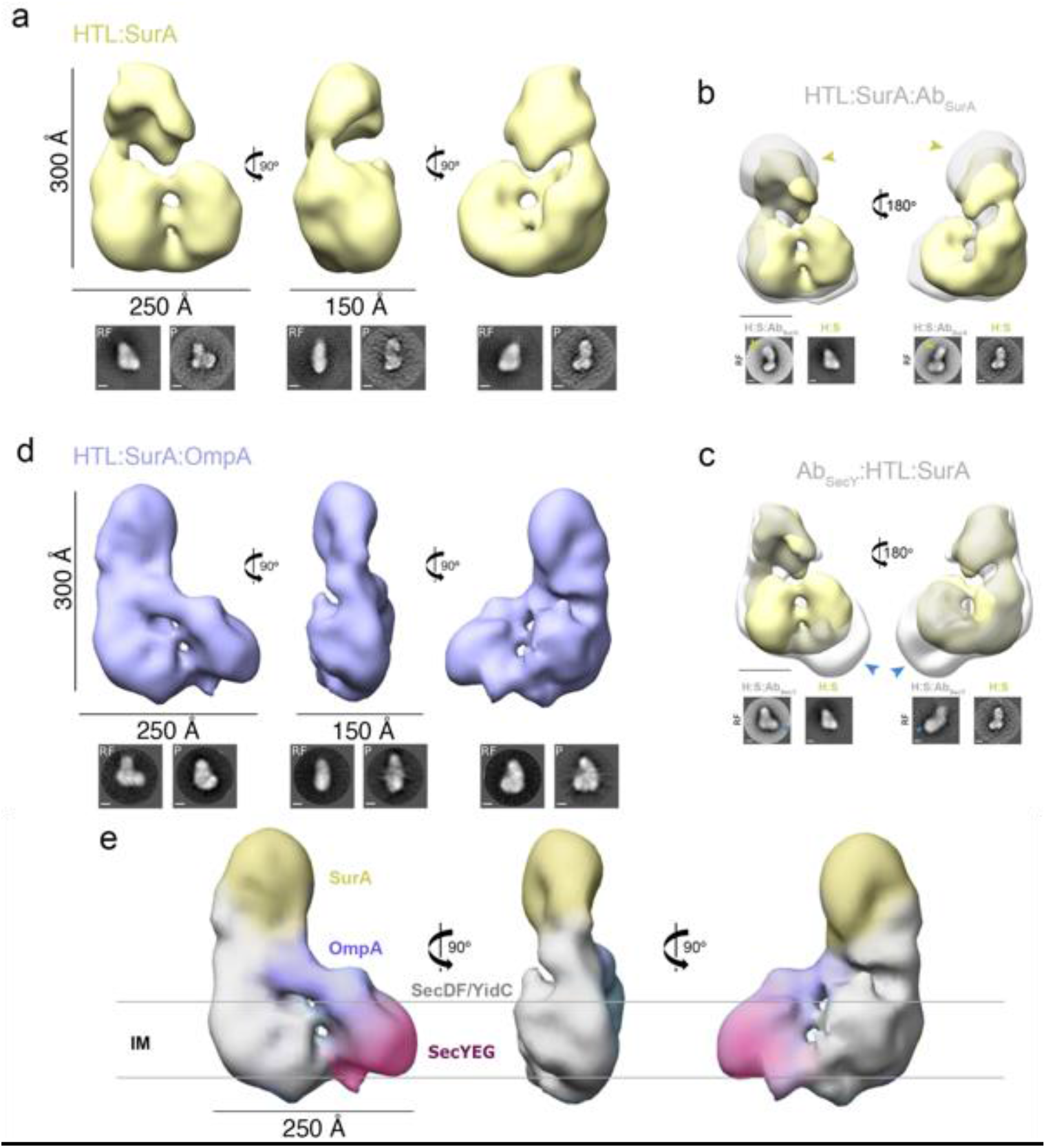
Negative-stain analysis of HTL:SurA and HTL:SurA-OmpA complexes: For a, b, c and d top panel shows views of 3D reconstructions, and bottom shows reference-free (RF) class averages and projections (P) of the final model, shown in the same orientations as the top. Scale bars are 100 Å, unless stated otherwise. See supplementary Figures 2 and 4 for image processing details. a) Orthogonal views of the HTL:SurA complex 3D reconstruction (41.9 Å resolution). b) Opposing side views of the HTL:SurA complex (yellow) superimposed on the immuno-complex HTL:SurA:Ab_SurA_ complex (grey). The antibody (yellow arrows) is shown bound to the mass assigned to SurA. c) Opposing side views of the HTL:SurA complex (yellow) superimposed on the immuno-complex Ab_SecY_ :HTL:SurA complex (grey). The antibody (blue arrows) is shown bound to the mass assigned to SecYEG. d) Orthogonal side views of the HTL:SurA-OmpA complex 3D reconstruction (37.5 Å resolution). e) Assigned map of the HTL:SurA-OmpA complex in the three representative orthogonal views, with components coloured accordingly.

The low-resolution analysis by negative stain visualised a large object measuring ~300 Å × 200 Å × 150 Å, exceeding the expected dimensions of HTL alone, even in its ‘open’ state (~200 Å × 200 Å × 150 Å) [23]. The larger size implicates the presence of SurA within the density, consistent with the mass spectrometry. However, the resolution was insufficient to localise the individual constituents of the stabilised HTL-SurA complex.

### SURA INTERACTS WITH PERIPLASMIC REGIONS OF SECDF-YIDC

In order to locate SurA within the complex with HTL, the density gradients were repeated following the addition of antibodies raised against either SecY (monoclonal) or SurA (polyclonal) (supplementary Figure S3a, b). Similarly to the previous sample, negative stain EM samples were prepared (supplementary Figure S3c, d); reconstructions of the cross-linked samples containing the antibodies were then overlaid with the reconstruction of HTL:SurA. Additional densities (yellow and blue arrows in Figure 2b and c) identify the respective antibodies, and thereby reveal the locations of SurA and SecY in the complex. The SecY monoclonal antibody recognises the cytosolic face of the protein-channel complex [24], which is located at the opposite side to the SurA, as expected on the periplasmic surface. Presumably then, SurA is associated with constituents of the HTL: SecDF and YidC, which possess suitable large periplasmic domains capable of forming interactions with hydrophobic substrates [25–27], whereas SecYEG does not [4].

### SURA IS IDEALLY SITUATED ON THE HTL TO COLLECT PROTEINS EMERGING FROM THE EXIT SITE OF SECYEG

The SurA chaperone can be purified either alone or together with an OMP substrate, enabling the analysis as described above. Similarly, to HTL:SurA the high molecular weight complex containing also OmpA (Figure 1d –asterisk) was analysed by negative stain electron microscopy (supplementary Figure S4) to obtain a low-resolution structural information of the substrate engaged assembly (HTL:SurA:OmpA; Figure 2d). When comparing the structures of the occupied and vacant chaperone-HTL complex, an extra density is apparent in the former which can be assigned to the OmpA substrate (Figure 2e). The positioning of OmpA is in contact with periplasmic surface of SecYEG, localised by the monoclonal antibody against SecY on the cytosolic side of the channel (Figure 2c). At this position it is roughly adjacent to the protein-channel exit site and in contact with SurA, possibly representing a late stage inner-membrane translocation intermediate. Presumably, SurA binds the translocon with its substrate binding site facing towards the protein exit channel of SecYEG ready for polypeptide capture.

### OMPA STABILISES INTERACTIONS BETWEEN HTL AND THE SURA CHAPERONE

The processing workflows of the negative stain EM data reveals that there is a large degree of flexibility between SurA and the HTL within the HTL:SurA complex (supplementary Figures S2), which is much reduced in the presence of OmpA (supplementary Figures S4). This suggests the OmpA somehow stabilises the chaperone-translocon interaction. To further explore this affect we employed size-exclusion chromatography, in the presence of a low concentration of an amide crosslinker (to stabilise the low-affinity interactions). This was conducted for the translocon alone, or following addition of unoccupied or substrate-engaged SurA (Figure 3a). In the presence of the OMP substrate there is a large peak at a higher apparent molecular weight, compared to the translocon alone, or empty chaperone bound complex (Figure 3a, asterisk). Subsequent, analysis of this fraction by SDS-PAGE and negative stain EM (Figure 3b, c) confirms the presence of a cross-linked complex of the similar dimensions and shape to the complexes isolated by density gradient centrifugation (Figure 2). The large shift of the OmpA engaged chaperone-translocon substrate is indeed consistent with change in the dynamic properties of the complex, compared to the unoccupied complex. This is in keeping with an expected response to the presence of OmpA and known changes of inter-domain dynamics of SurA upon interaction with its clients [13].

**Figure 3:**
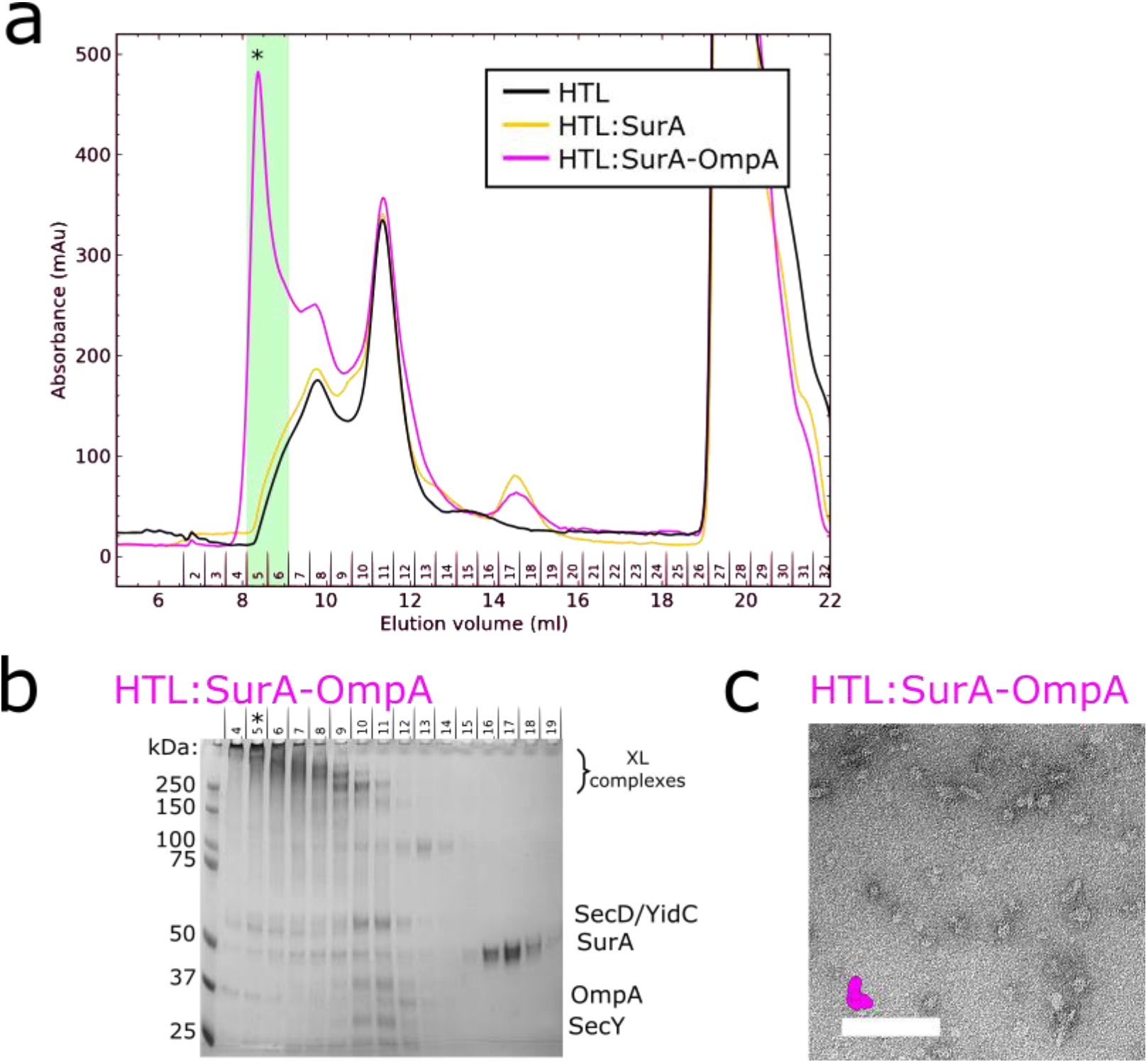
HTL:SurA-OmpA complex isolation by size-exclusion chromatography: a) Size exclusion chromatograms for crosslinked samples containing purified HTL alone (black) or pre-incubated with SurA (yellow) or SurA-OmpA complexes (magenta). b) SDS-PAGE analysis of fractions corresponding to the chromatogram in (a). c) Negative stain micrographs prepared from fractions highlighted in (b) with an asterisk. Scale bars represent 1000 Å. For indication of correct complex size, the reconstruction from the GraFix experiment for HTL:SurA-OmpA is depicted in scale in magenta.

### LOW-RESOLUTION CRYO-EM STRUCTURE ANALYSIS REVEALS COMPLEX DISORDER

Cryo-EM analysis of the high-molecular weight peak from the HTL:SurA:OmpA size exclusion chromatography (Figure 3a, asterisk; supplementary Figure S5) resulted in the reconstruction shown in Figure 4. By comparison to the negative stain structures (Figure 2) we can assign the locations of SurA and the constituents of the HTL. The low-resolution attainable is an unfortunate consequence of the known inherent flexibility of the HTL [23,23] and SurA (see below), required for their function. The associated OmpA is presumably also very flexible and not well defined at this resolution (Figure 4c, arrow). The poorly resolved unfolded OmpA likely contributes to the noise of the reconstruction, particularly between the chaperone and membrane protein complex, which was evident during processing (supplementary Figure S5).

**Figure 4:**
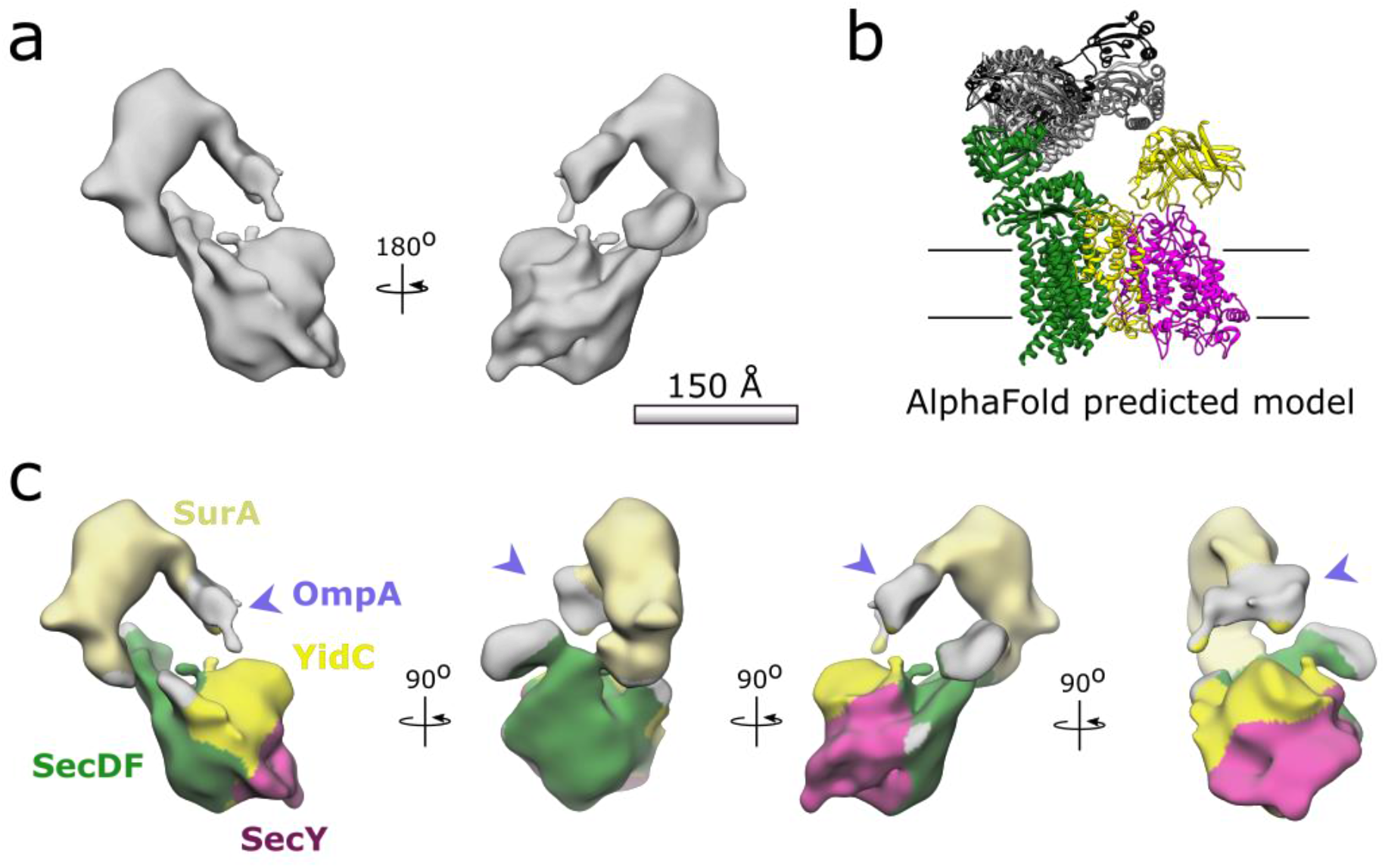
Low resolution cryo-EM of HTL:SurA:OmpA: a) 22 Å resolution reconstruction processed using Relion, see supplementary figure 5 for image processing details. b) The top five ranked AlphaFold predictions for the interactions between *E.coli* SurA (different shades of grey) and *Thermus thermophilus* SecDF (green). YidC (yellow) and SecYEG (magenta) have been added according relative positions in the published HTL structure PDB:5MG3 [31]. c) Locations for SurA (light yellow) and SecY (magenta) are both consistent with negative stain analysis in Figure 2 and the Alphafold prediction in (b). As in (b) locations of SecDF (green) and YidC (yellow) have been mapped relative to SecYEG according to PDB:5MG3 [31]. The purple arrow indicates extra density likely representative of OmpA.

It is now established that SurA is highly dynamic [13–16] with several different conformations for OmpA binding [16]. With this in mind, it is unlikely that there is a single conformation for the HTL:SurA:OmpA complex, which is problematic for sample preparation and image processing (supplementary Figure S5). Despite the low resolution achievable the negative stain and cryo-EM have been able to confirm an important interaction of a periplasmic chaperone with the Sec translocon for collection of nascent envelope proteins.

### ALPHAFOLD PREDICTS AN INTERACTION BETWEEN SECDF AND SURA

Through AlphaFold2 [29] we built a model for the most likely interaction between the HTL and the chaperone (Figure 4b) and used this to inform our prediction of the subunit organisation depicted in Figure 4c. Each of the core components of HTL was run through the structural prediction software in the presence of the SurA chaperone.

The SurA interactions with YidC and SecYEG (supplementary Figure 6e, f) were varied but consisted mostly with cytosolic or membrane regions of the inner membrane proteins hence unfeasible within the context of the membrane envelope. However, all five top ranked models for interactions between SurA and SecDF showed SurA bound with the periplasmic P1-head domain of SecDF in an extended I-form conformation (supplementary Figure 6 b, d) (observed in [30]). The top ranking prediction of the SecDF:SurA interaction was then contextualised within the HTL using known subunit organisation [31] to produce the model in Figure 4b. This has striking similarities to our observed densities from both negative stain and cryo-EM. Additionally the SurA in the five models predicted by AlphaFold presumes a variety of conformations consistent with our observed structural heterogeneity.

Importantly however, in the predicted models the contact between the SurA chaperones and the SecDF did not alter. Each model showed an interaction between the P1 and P2 parvulin-like PPIase domains of SurA and the periplasmic P1 head domain of SecDF. This is of particular interest as the P1 domain in SecDF is thought to have polypeptide binding capabilities, and moves in response to proton transport [30].

Additionally, previous AlphaFold models for SecDF alone (entry AF-Q5SKE6-F1 on the AlphaFold Protein Structure Database) turn out with the inactive super-F form [32]. However, in the presence of SurA AlphaFold predicts the active, extended form of SecDF. This suggests a preferred conformation of SecDF for SurA binding, which could be regulated during proton transport driven by the PMF.

## DISCUSSION

From the findings shown in this paper we can now present an updated model for OMP targeting (Figure 5). We have established that SurA can interact with the large periplasmic domains of the HTL; presumably with either SecDF – already known to facilitate protein secretion [33,34], or with YidC, or possibly both. Recently, we have proposed that SecDF facilitates the onward passage of OMPs to the outer-membrane through a direct interaction with BAM [23]. We also speculated that the periplasmic chaperones SurA, Skp, PpiD and YfgM are also involved. The results described here suggest that this is likely to be true, at least in the case of the former candidate. The disposition of SurA is ideal for welcoming proteins emerging through SecYEG. The dynamic properties of SurA and the periplasmic regions of SecDF, noted here and elsewhere [13–16,23,25,27,28,30,31], are presumably exploited during the passage of proteins from the Sec-machinery to BAM.

**Figure 5:**
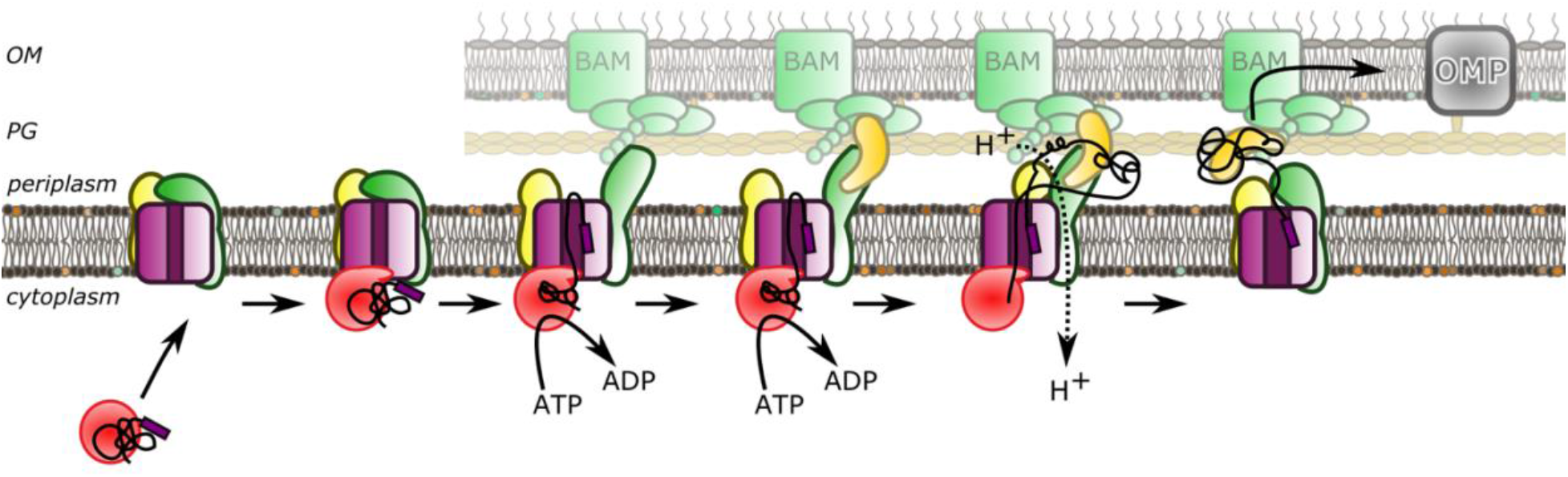
A model for OMP secretion across the bacterial envelope: OMPs are targeted to the inner membrane via an association between SecA (red), and the holo-translocon (SecY;magenta, SecDF;dark green, YidC;yellow). Translocation across the inner membrane is driven by ATP hydrolysis by SecA. SurA (gold) interacts with the OMP during the translocation process through interactions with YidC and/or SecDF (yellow and green) and the OMP. OMP targeting to the outer membrane insertion machinery BAM (bright green) is assisted through interactions between the HTL and BAM and/or between SurA and BAM. Outer membrane insertion can then occur through the BAM machinery.

In the updated model, while pre-protein translocation is still underway – driven through SecYEG by the motor ATPase SecA – the emerging protein immediately checks in with SurA (or perhaps another suitable chaperone), which prevents misfolding or aggregation prior to substrate release (Figure 5). Consistent with this, *surA* mutants have previously been shown to affect OMP release from the Sec-machinery [19]. Following release from the inner-membrane SurA and its client are presumably then free to move in order to facilitate OMP transfer to the connected BAM for outer-membrane insertion. Potentially, the free energy available in the PMF, driving conformational changes of the connected domains of SecDF [30], and possibly YidC, drive this process in the desired direction; *i.e*. a net flow of protein to the outer-membrane (Figure 5). This could be achieved *via* interactions of their periplasmic regions with the chaperone (shown here), or with the OMP directly. The respective domains of SecDF and YidC both contain clefts suitable for binding hydrophobic stretches of polypeptides typical of OMPs [26,30].

It seems likely that the associated chaperone maintains the OMPs in an unfolded aggregation free state prior to their release to BAM. Interestingly, the recently described Sec-BAM super-complex contains a central cavity, between the two membranes, large enough to accommodate small chaperones like SurA [23]. Given the known interaction of SurA with both HTL and BAM, it is possible that it could be very intimately associated within the *inner sanctum* of a trans-envelope super-complex. Alternatively, SurA could form part of the walls or gate of the periplasmic chamber. A protected environment like this would of course help prevent aggregation, proteolysis and provide an easier route through the periplasm and PG layer. In addition, the open nature of this cavity would also allow lateral escape of periplasmic proteins for folding, and for the release of misfolded proteins for degradation. In this respect, SurA may also help distinguish globular proteins destined for the periplasm, from β-barrelled outer-membrane proteins, and thereby regulate their lateral release and folding.

In addition to the role in outer-membrane targeting, chaperone binding of the OMP substrate during the translocation process could also help drive the translocation process itself. One of the models for SecA-dependent transport through the inner membrane suggests that the SecY channel facilitates a Brownian ratchet mechanism for biased diffusion through the channel [35,36]. Here backsliding of positively charged and bulky residues is prohibited by channel closure following retraction of the 2-helix finger of SecA from SecY. Chaperone binding to the unfolded protein substrate during translocation would inhibit backward diffusion of the substrate through the SecY thereby contributing to the ratchet and promoting translocation. Protein folding in the periplasmic cavity of SecY has been shown to have a similar effect [37].

Together, these findings present a novel interaction which improves our understanding of protein secretion in Gram-negative bacteria. The HTL-SurA complex is particularly important as it is unique to bacteria and therefore presents a promising target for therapeutic inhibition by antibiotic development.

## METHODS

### Chemical reagents

*Escherichia coli* polar lipid and cardiolipin (CL) were purchased from Avanti, and were prepared at 10 mg/ml in 50 mM triethanolamine, pH ~ 7.5, 50 mM KCl.

### Cell strains

Chemically competent C43 cells were used for expression of HTL. These were cultured from home-made stocks of cells gifted by Prof. Sir John Walker (MRC, Mitochondrial biology unit, Cambridge). Expression of all soluble proteins used chemically competent home-made stocks of E.coli strain BL21(DE3) originally sourced commercially (NEB).

### Protein expression

#### HTL

Chemically competent C43 cells were used for expression of HTL [38]. These were cultured from home-made stocks of cells gifted by Prof. Sir John Walker (MRC, Mitochondrial biology unit, Cambridge). HTL was purified as described previously [21,39,40] in TS_130_G buffer (20 mM Tris-HCl^pH≈8^, 130 mM NaCl, 10% (v/v) Glycerol) for density gradient experiments and with Tris substitution for 50 mM HEPES^pH≈8^ in size exclusion experiments (HS130G buffer: 50 mM HEPES^pH≈8^, 130 mM NaCl, 10% (v/v) Glycerol) [21,39,40]. Throughout later stages of purification both buffers were supplemented with 0.02% cardiolipin (CL) as described previously [23].

#### SurA-OmpA

*pET28b-His-surA* and *pET11a-ompA* [41] were a gifts from Prof. Sheena Radford (The Astbury Centre for Structural Molecular Biology, University of Leeds, UK). Expression of all SurA and SurA-OmpA proteins was carried out using chemically competent home-made stocks of E.coli strain BL21(DE3) originally sourced commercially (NEB). Both proteins were over-produced separately in 1 L of cultures as described previously [42]. Both were harvested by centrifugation, and resuspended in TS_130_G (20 mL 20 mM Tris pH 8.0, 130 mM NaCl, 10% (v/v) glycerol) in the presence of c*O*mplete protein inhibitor cocktail (Roche). Both were then lysed in a cell disruptor (Constant Systems Ltd.) and the resulting samples were clarified by centrifugation in an SS34 rotor (Sorvall) at 27,000 ×*g*, 4°C for 20 minutes. For OmpA, the supernatant was discarded and the pellet resuspended in 20 mL TS_130_G + 6 M urea. The OmpA pellet was diluted to 80 mL with 6 M urea and mixed with the SurA supernatant to give a final urea concentration of 4.8 M. The urea was removed by dialysing in 2 L TS_130_G for 6 hours at room temperature, then dialysing overnight at 4°C in 5 L fresh TS_130_G. The sample was centrifuged again 27,000 ×*g*, 4°C for 20 minutes and the supernatant loaded onto a 5 mL HisTrap HP column (Cytiva) equilibrated with a low salt buffer, TS_50_G (20 mL 20 mM Tris pH 8.0, 50 mM NaCl, 10% (v/v) glycerol). The column was washed with a low imidazole buffer (TS_50_G +20 mM imidazole), and bound proteins eluted with a high imidazole buffer (TS_50_G + 300 mM imidazole). The eluents were loaded onto a HiTrap Q HP column (Cytiva) equilibrated in the low salt buffer (TS_50_G) and free SurA was found in the unbound fraction. A linear gradient of 0.05 - 1 M NaCl was applied over 60 mL and following SDS-PAGE analysis, fractions containing SurA-OmpA were pooled.

For size exclusion experiments, purification was repeated under identical conditions with Tris substituted with 50 mM HEPES during the purification (HS_20_G: 50 mM HEPES ^pH≈8^, 20 mM NaCl, 10 % glycerol). The flow-through from HiTrap Q HP column (Cytiva) contained the SurA-OmpA under these conditions and so this was concentrated and to <500 μl and loaded to a S200 30/150 SEC column (Sigma). Fractions from the peaks corresponding to SurA-OmpA complexes were pooled and concentrated.

### Binding assay between HTL and SurA or SurA:OmpA

HTL (10 μM) was incubated with purified SurA or SurA:OmpA at 1:2 molar ratio in TS_130_G buffer (20 mM Tris-HCl^pH≈8^, 130 mM NaCl, 10% (v/v) Glycerol) supplemented with 0.01% DDM-CL for 30 min at 30 °C. Individual proteins were subjected to the same procedure. Aliquots were then loaded to a native electrophoresis using NativePAGE Bis-Tris 4-16% (Invitrogen) gels run at 140 V for 2 hours at room temperature. Protein complex bands separated in the native electrophoresis were excised, incubated for 10 min in SDS-PAGE sample buffer, applied to the top of a SDS-PAGE gel NuPAGE 4-12%, 1.5mm wells (Invitrogen) and run at 140 V for 90 minutes. Gel bands of the individual proteins were excised at the same height as the protein complexes as migration controls. After electrophoresis, gels were silver stained with SilverQuest™, (Invitrogen) following manufacturers recommendations.

### Density gradient ultracentrifugation

Glycerol gradients for gradient fixation (GraFix) as described in [22] were prepared in a Sw60Ti ultracentrifugation tube (Beckmann) with 26 layers of 150 μL per glycerol concentration. The different concentrations were achieved by mixing a 0% glycerol gradient buffer (20mM HEPES^pH≈8^, 250mM NaCl, 0.03% DDM-CL, with a 60% glycerol buffer (20 mM HEPES^pH≈8^, 250 mM NaCl, 0.03% DDM-CL, 60% glycerol) to achieve a gradient from 20-40% glycerol without or alongside a glutaraldehyde gradient of 0-0.15%.

Different combinations of HTL alone or with either SurA or SurA:OmpA complexes were diluted in the 0% glycerol gradient buffer (20 mM HEPES, 250 mM NaCl, 0.03% DDM-CL) to a total volume of 150 μL with protein concentrations of 5 μM each. These were incubated at 30 °C for 40 minutes followed by 5 minutes on ice. Protein mixtures were carefully floated on top of prepared gradients and centrifuged in a Sw60Ti rotor (Beckmann) for 16 hours at 134,000 × g and 4 °C. Each layer was manually extracted into tubes analysed by SDS-PAGE with silver staining using the fast staining protocol within SilverQuest™ Silver staining kit (# LC6070 Novex).

For HTL:SurA:Ab_SurA_ and Ab_SecY_:HTL:SurA complexes, 5 μM of HTL and SurA were incubated in binding buffer at 30°C for 30 minutes with shaking in a total volume of 150 μl. SecY antibody was added at a concentration of 1 in 7.5 and incubated at 30°C for 15 minutes with shaking in a total volume of 150 μl.

### Size exclusion chromatography (SEC)

10 μM of HTL and/or SurA/SurA-OmpA were mixed in 150 μL reaction volume in buffer containing 50 mM HEPES^pH≈8^, 130 mM NaCl, 10% glycerol, 0.01% DDM-CL and 1 μg/ml pepstatin A. Reactions were incubated at 25°C for 30 minutes with shaking. Bis- (Sulfosuccinimidyl) suberate (BS3, Thermo Fisher Scientific #21580) was added to a concentration of 3.2 mM and further incubated for 40 minutes at 25°C with shaking. Finally 50 mM ammonium chloride was added to quench the crosslinking and this was incubated at 25°C for a further 15 minutes. Samples were then loaded onto a S200 10/300 column (Sigma) equilibrated in buffer containing 50 mM HEPES^pH≈8^, 130 mM NaCl, 0.01% DDM-CL (no glycerol). Fractions were analysed through SDS-PAGE with Coomassie staining.

### Negative Stain Electron Microscopy

#### Sample preparation

Negative stain EM was conducted using C300Cu100 (agar scientific) carbon coated copper grids glow discharged for 15s. 5 μl of a suitably diluted protein sample was applied to the carbon side of the grid and incubated for ~1 minute. Grids were blotted, washed with water, re-blotted and finally stained using a 5 μl drop of 2% uranyl acetate for a further 1 minute. This was blotted a final time and grids were stored and logged. All further analysis was done using either a Tecnai 12 microscope at the Living Systems Institute, University of Exeter, or the Tecnai 12 and Tecnai 20 microscopes in the Wolfson Bioimaging Suite at the University of Bristol (see below):

##### GraFix purified HTL-SurA and HTL-SurA-OmpA

Tecnai 12 (Thermo Fisher Scientific) with a One View Camera (Gatan) at a digital magnification of 59,400 × and a sampling resolution of 2.1 Å per pixel (EM facility, University of Exeter)

##### SEC purified HTL-SurA-OmpA

Tecnai 20 (Thermo Fisher Scientific) with Ceta 4k × 4k CCD camera (Thermo Fisher Scientific) at a digital magnification of 50,000 and a sampling resolution of 2.02 Å per pixel (Wolfson Bioimaging facility, University of Bristol)

###### Processing

All image processing for negative stain EM was carried out in EM software framework Scipionv2.0 [43] using University of Bristol High Performance Computing specialist cluster BlueCryo. Micrographs were imported and manually picked particles provided the input toward automatic picking through the XMIPP3 package [44,45]. Particle extraction used Relion v2.1 package [46,47] and reference-free class averages grouped particles using Relion 2D classification (v3.0). The low quality classes from this output were removed and a second round of 2D classification was conducted. Similar 2D classification software XMIPP-CL2D was also carried out in parallel [48]. The outputs of these could be used to generate 3D models using EMAN v2.12 software [47,49]. Many consecutive rounds of 3D classification in Relion (v3.0) [47] were used to filter out particles and create a homogeneous set for the most detailed structure possible. This was carried out using Relion 3D-classification. 3D auto-refinement was also conducted using Relion (v3.0). Resolution estimates through Relion used 0.143 correlation coefficient criteria [50,51].

### Cryo-Electron Microscopy

4 μL of marked fractions from size exclusion chromatography (asterisk, Figure 3) diluted 40 times (in 50mM HEPES^pH≈8^, 130mM NaCl, 0.01% DDM-CL) were loaded on freshly prepared graphene oxide coated grids (prepared using methods described in [52]) and plunge frozen into liquid ethane using a Vitrobot Mark IV unit (Thermo Fisher Scientific) with wait time of 2 seconds after sample application and blotting time of 2 seconds. 1081 movies acquired on a TALOS Arctica (Thermo Fisher Scientific) microscope with K2 direct electron detector (Gatan) in linear mode (at GW4 facility, University of Bristol).

Patch motion correction was carried out using cryoSPARC v2.15 + 200728 4.4 within the cryoSPARC web interface [53–56]. The remainder of the processing was conducted within the Scipion v2.0 framework [43] and included use of: CTFFIND4 [57,58], Xmipp v3.0 [45,45] and Relion v3.0 [49,47,51,60].

### AlphaFold

AlphaFoldv2 [29] was carried out within Google Colab [61]. *E.coli* amino acid sequences were used for SurA (P0ABZ6) and YidC (P25714). To reduce complexity the single chain SecDF from *Thermus thermophilus* (Q5SKE6) was used. *T.thermophilus* SecDF is also used in a number of structural studies and has high levels of conservation to SecDF in *E.coli* [25,32].

### Graphical analysis

Molecular graphics and analyses was performed with UCSF Chimera v1.15, developed by the Resource for Biocomputing, Visualization, and Informatics at the University of California, San Francisco [62]. Any map segmentation was done using the segger tool within this software [63,64]. As stated, molecular ‘docking’ was conducted by hand. The coloured map in figure 4c used Chimera ‘Color Zone’ following the AlphaFold model to inform the docking of HTL (PDB:5mg3) and SurA (PDB:1m5y) into the cryo-EM density.

## Acknowledgments

We gratefully acknowledge access and support of the Wolfson Bioimaging Facility and the GW4 Facility for High-Resolution Electron Cryo-Microscopy, with particular thanks to Dr Ufuk Borucu. Thanks to Kate Heesom and the Proteomics Facility at the Faculty of Life Sciences (University of Bristol) for the mass-spectroscopy analysis. We are also grateful to Mathew McLaren for support with electron microscopy at the University of Exeter.

## Author contribution

LT designed and conducted experiments, assisted by SA; LT and IC wrote the manuscript; IC secured funding and led the project; VG and BD provided resources and equipment for electron microscopy imaging in Exeter.

## Declaration

the authors declare no competing interests.

## Data and materials availability

All data are available in the main text or the supplementary materials.

---

**Supplementary Figure 1:**
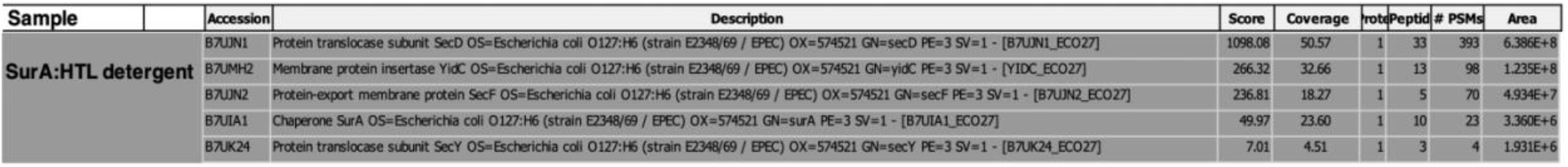
Mass-spectrometry analysis of the HTL:SurA complex purified by density gradient centrifugation: The fraction denoted with an asterisk in Figure 1c was taken for this analysis. This confirms the presence of SecD, YidC, SecF, SurA and SecY within the sample.

**Supplementary Figure 2:**
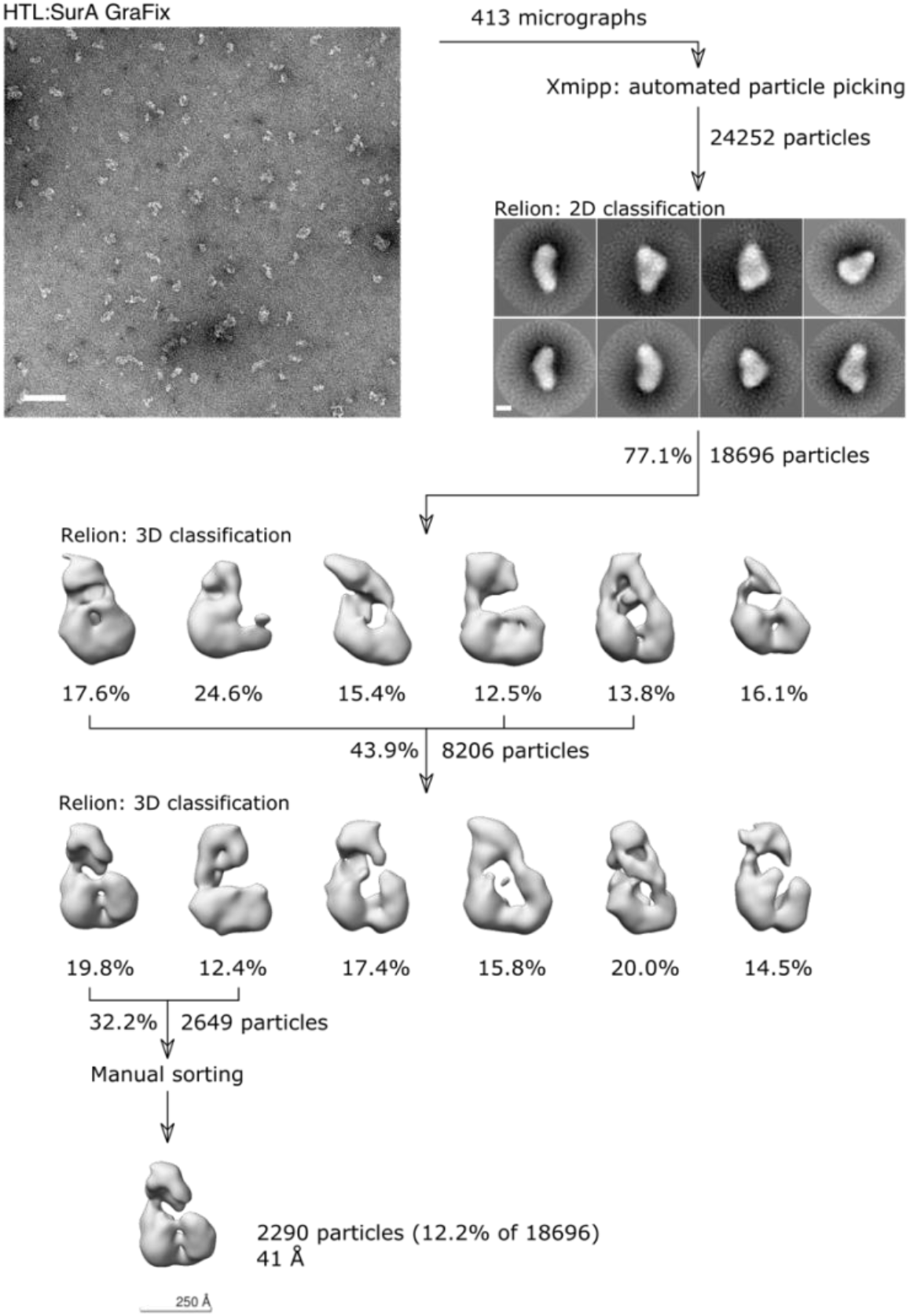
Negative-stain HTL:SurA processing pipeline: Images of 413 micrographs yielded 24,252 particles following picking, of which a subset of 18,696 quality particles were isolated through 2D classification. Following Relion 3D classification, classes revealed a variety of conformations and particle qualities. Three of the most similar classes were chosen with 8,206 particles. A second classification followed by a manual particle sort prior to relion autorefine resulted in a low-resolution structure of 41 Å containing only 12.2% of the original particles. Scale bars for the micrograph (top left) and reference free 2D class averages (top right) represent 1000 Å and 100 Å respectively.

**Supplementary Figure 3:**
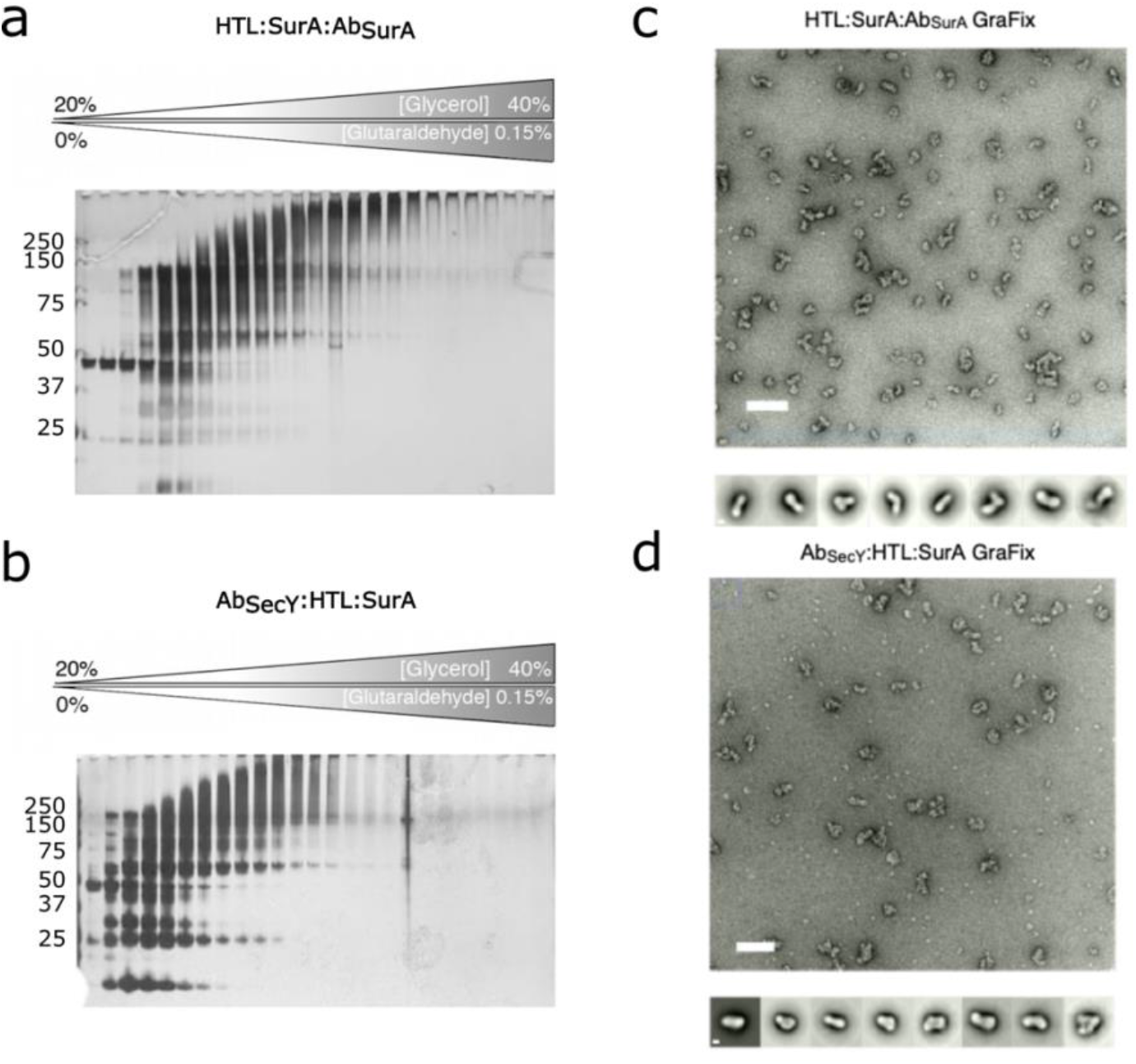
Density gradient centrifugation of the translocon and chaperones alongside SurA and SecY antibodies: a,b) Silver stained SDS-PAGE analysis of glycerol fractions following density gradient centrifugation of the translocon with SurA chaperone repeated with polyclonal SurA antibody (a) or a monoclonal SecY antibody (b). c,d) Representative negative stain micrograph and reference free 2D class averages of a single fraction of the density gradient centrifugation with SurA antibody (a,c) or SecY antibody (b,e). Scale bars for the micrograph and reference free 2D class averages represent 1000 Å and 100 Å respectively.

**Supplementary Figure 4:**
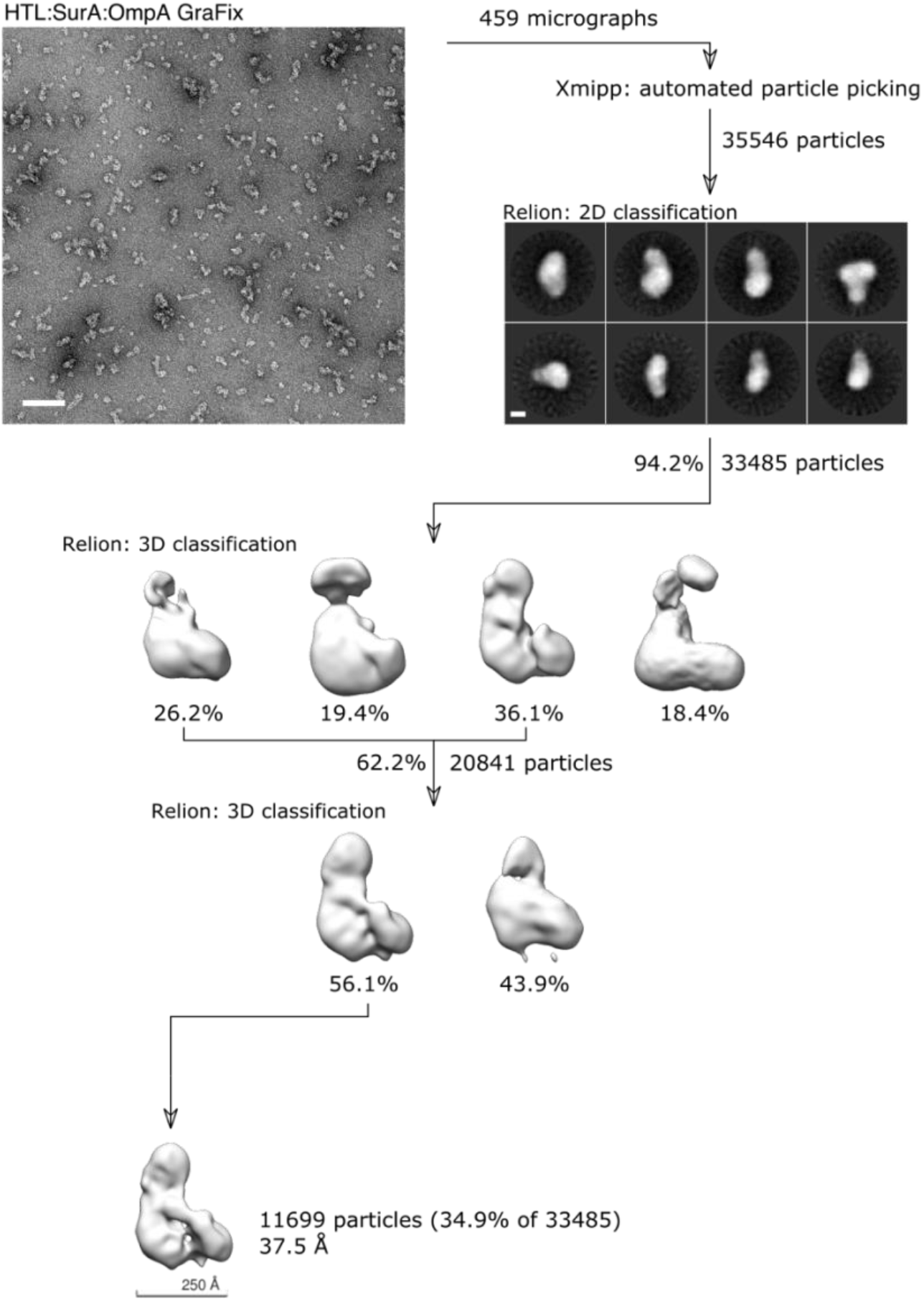
Negative-stain HTL:SurA-OmpA processing pipeline: Images of 459 micrographs yielded 35,546 particles following picking, of which a subset of 33,485 quality particles were isolated through reference-free 2D classification. Following Relion 3D classification, classes relion autorefine resulted in a low-resolution structure of 37.5 Å containing 37.5% of the original particles. Scale bars for the micrograph (top left) and reference free 2D class averages (top right) represent 1000 Å and 100 Å respectively.

**Supplementary Figure 5:**
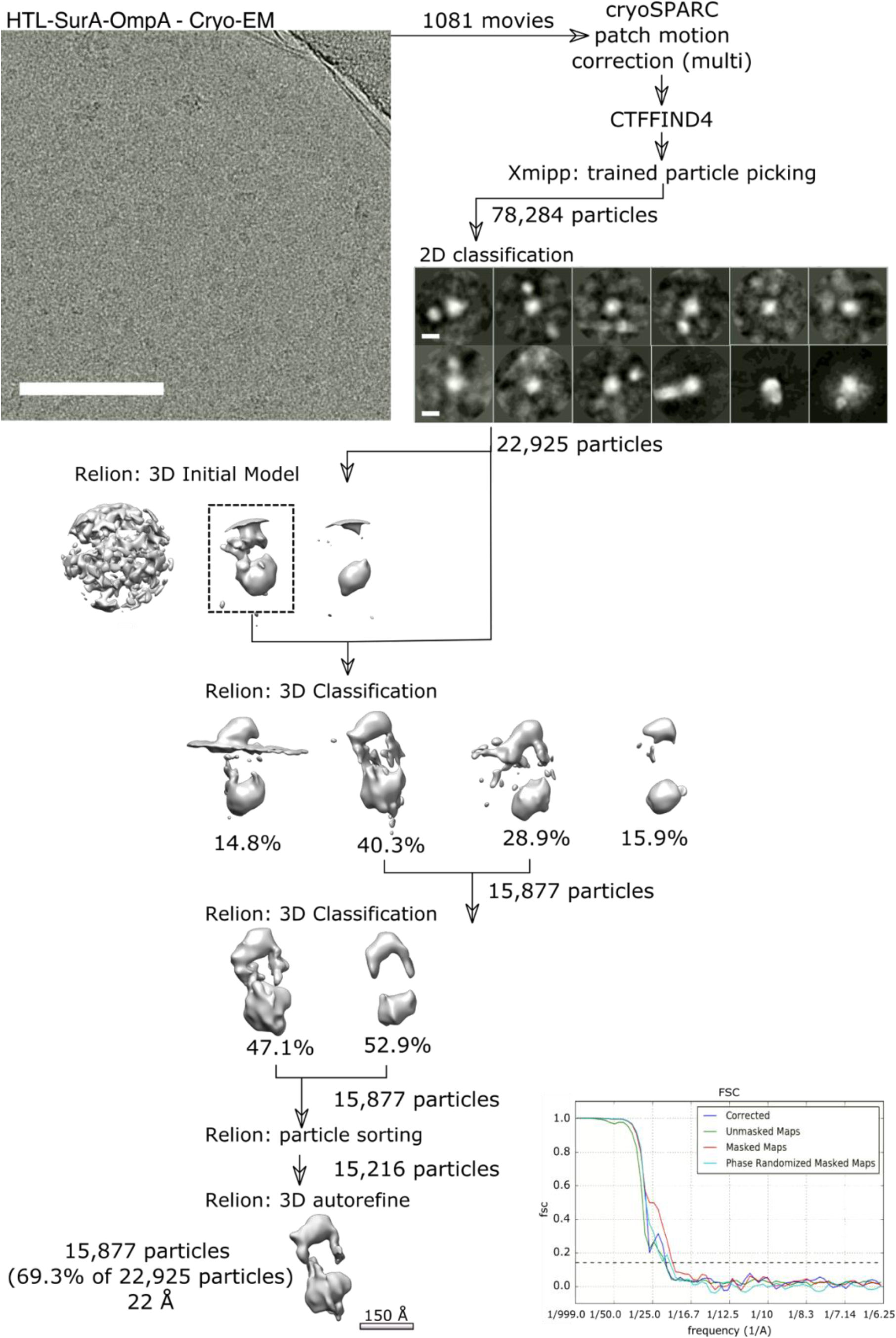
Cryo-EM data processing of HTL:SurA:OmpA: cryoSPARC patch motion correction aligned frames from 1081 movies, at this point micrographs were exported from processing in Scipion v2.0. Xmipp trained particle picking selected 78,284 particles, of which only 22,925 appeared the right size or had two clear densities (corresponding to SurA and HTL) for downstream processing. These were used to generate a reference-free initial model to inform two consecutive rounds of 3D classification. Following particle sorting a final run of relion 3D autorefine generated a map using 22,925 particles, or 69% of the particles selected following 2D classification. Resolution estimates through Relion gave a final resolution of 22 Å using the 0.143 correlation coefficient criteria, the plot of the Fourier shell correlation is shown in the bottom right. Scale bars for the micrograph (top left) and reference free 2D class averages (top right) represent 1000 Å and 100 Å respectively.

**Supplementary Figure 6:**
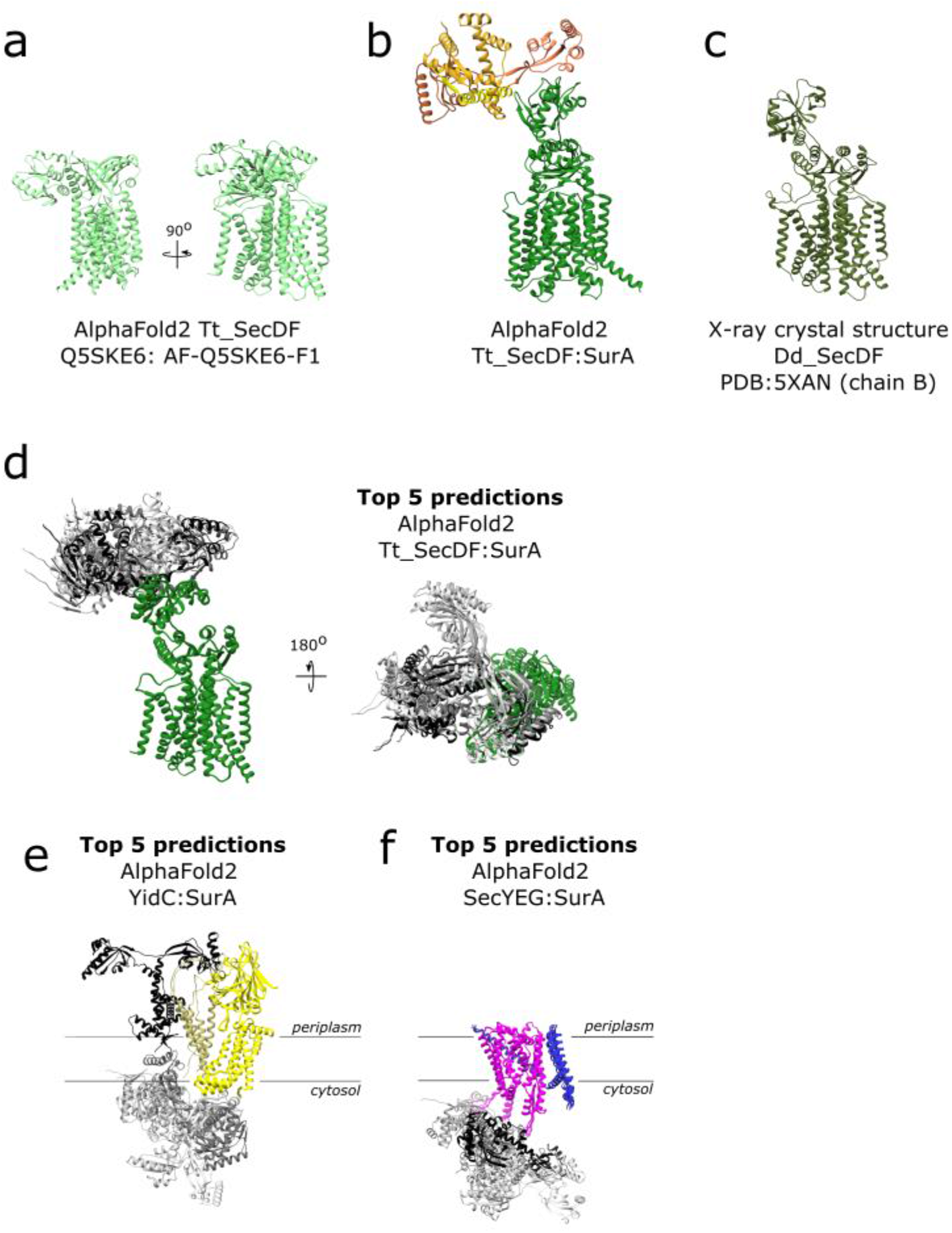
AlphaFold predictions for the HTL chaperone interaction: a) AlphaFold predicted structure for SecDF from *Thermus thermophilus* (result number AF-Q5SKE6-F1 on AlphaFold protein structure database). b) The top ranking result for AlphaFold2 run with SecDF from *Thermus thermophilus* and SurA from *E.coli*. c) X-ray crystal structure of SecDF in I-form conformation from *Deinococcus radiodurans R1* PDB:5XANb [30]. d) The top 5 predictions for AlphaFold2 run with SecDF (green) from *Thermus thermophilus* and SurA from *E.coli* (in different shades of grey) from the side and viewed from the top. e) The top 5 predictions for AlphaFold2 run with YidC (yellow) and SurA (in different shades of grey) from *E.coli*. f) The top 5 predictions for AlphaFold2 run with SecY (magenta), SecE (blue) and SurA (in different shades of grey) from *E.coli*.

